# Quantitative LCMS for ceramides/dihydroceramides: pregnancy baseline biomarkers and potential metabolic messengers

**DOI:** 10.1101/2020.02.24.963462

**Authors:** Qianyang Huang, Shiying Hao, Xiaoming Yao, Jin You, Xiao Li, Donghai Lai, Chunle Han, James Schilling, Kuo Yuan Hwa, Sheeno Thyparambil, John Whitin, Harvey J. Cohen, Henry J. Chubb, Scott R. Ceresnak, Doff B. McElhinney, Gary M. Shaw, David K. Stevenson, Karl G. Sylvester, Xuefeng B. Ling

## Abstract

Ceramides and dihydroceramides are sphingolipids that present in abundance at the cellular membrane of eukaryotes. Although their metabolic dysregulation has been implicated in many diseases, our knowledge about circulating ceramide changes during the pregnancy remains limited. In this study, we present the development and validation of a high-throughput liquid chromatography-tandem mass spectrometric (LC/MS/MS) method for simultaneous quantification of 16 ceramides and 10 dihydroceramides in human serum within 5 mins by using stable isotope-labeled ceramides as internal standards (ISs). This method employs a protein precipitation method for high throughput sample preparation, reverse phase isocratic elusion for chromatographic separation, and Multiple Reaction Monitoring (MRM) for mass spectrometric detection. To qualify for clinical applications, our assay was validated against the FDA guidelines: the Lower Limit of Quantitation (LLOQ as low as 1 nM), linearity (R^2^>0.99), precision (Coefficient of Variation<15%), accuracy (Percent Error<15%), extraction recovery (>90%), stability (>85%), and carryover (<0.1%). With enhanced sensitivity and specificity from this method, we have, for the first time, determined the serological levels of ceramides and dihydroceramides to reveal unique temporal gestational patterns. Our approach could have value in providing insights into disorders of pregnancy.

## 1. Introduction

Ceramides and dihydroceramides are subfamilies of sphingolipids characterized by the attachment of the aliphatic moieties onto the sphingosine and sphinganine backbones through amide-linkage. The inherent heterogeneity of the fatty acid molecules structurally diversifies both families of chemicals by conferring the individual species with a distinctive combination of carbon number and saturation degree on the aliphatic moiety (Fig.1) [1]. Biosynthetically, ceramides are primarily generated by the de novo synthetic pathway from the endoplasmic reticulum (ER) via the collaborative action regulated by multiple enzymes in the presence of transient intermediates. Among the functional intermediates, dihydroceramides are known as the immediate precursors that can be directly converted to ceramides via oxidation catalyzed by dihydroceramide desaturase (DEGS1) [2]. Ceramides are present in abundance on the membrane of eukaryotic cells along with other integral membrane components, including trans-membrane proteins, sphingomyelins, cholesterols, and glycosphingolipids. These components collectively form characteristic regions known as glycolipid-enriched microdomains or lipid rafts on the eukaryotic cell membrane, which play fundamental roles in maintaining the integration and dynamic behavior of the lipid bilayer scaffold [3]. In addition to their importance for the structural folding of eukaryotic cell membranes, ceramides have also been recognized as bioactive signaling modulators for various essential cellular events, particularly cell proliferation, differentiation, migration, adhesion, and apoptosis [4].

**Fig.1.**
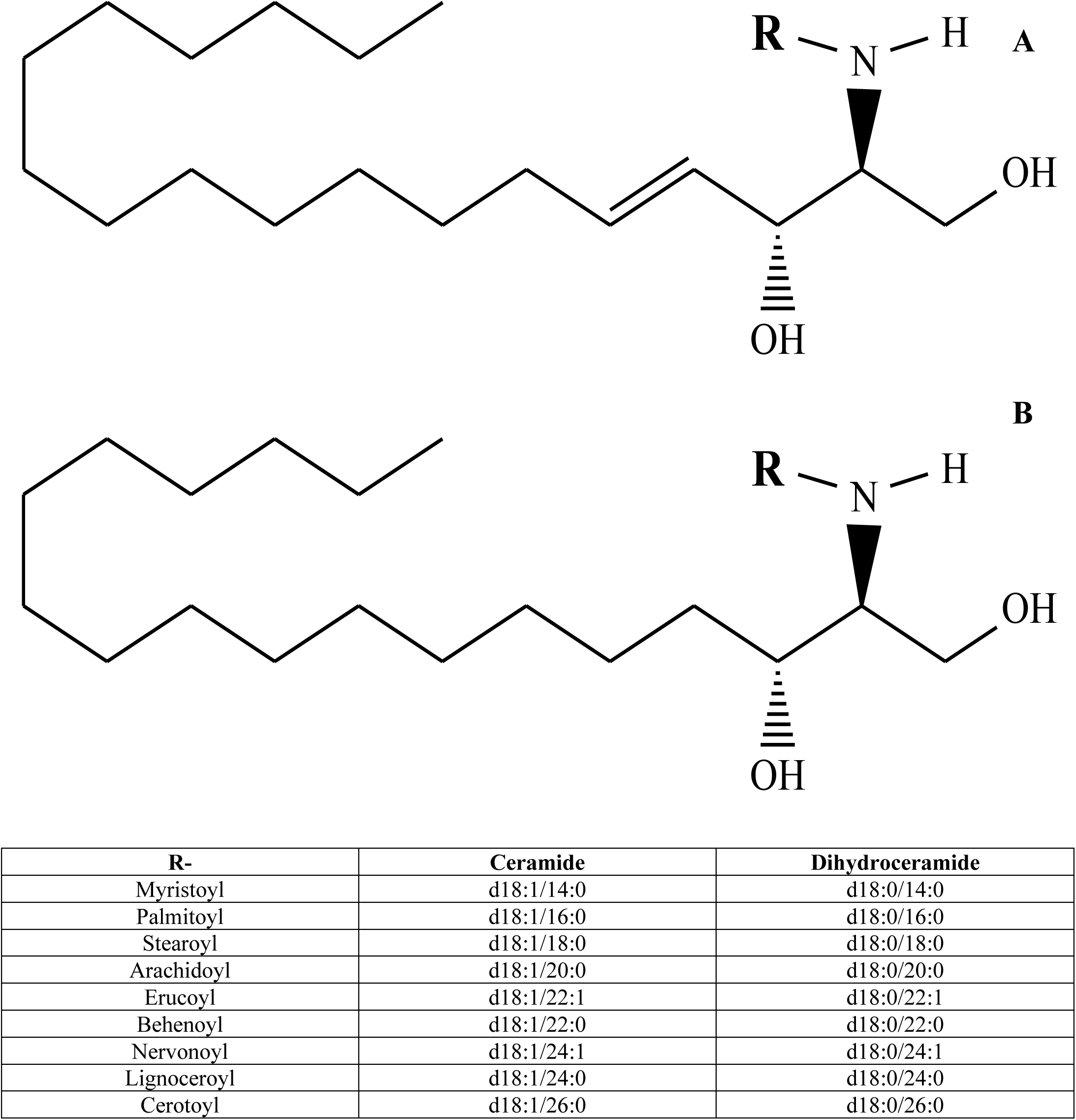
General structures and nomenclatures of ceramide (A) and dihydroceramide (B).

In view of their potential proapoptotic toxicity in normal eukaryotic cells, ceramide levels are dynamically regulated through metabolic influx from the catabolism of sphingomyelins and metabolic outflux through the degradation of ceramides into sphingosine-1-phosphate, a bioactive sphingolipid that is known to promote cell survival by counteracting the apoptotic effect induced by elevated ceramides. This balancing effect of ceramide metabolism creates an interactive rheostat for subcellular machinery to exert its regular physiological functions [5].

In contrast, alternation of this rheostat between cell apoptosis and survival has been reported to contribute to abnormal early development of the placenta, characterized by compromised proliferation, differentiation, and apoptosis of trophoblast cells. Such changes could lead to persistent placental hypoxia and oxidative stress due to insufficient uteroplacental circulation from the maternal spiral arteries and eventually to the pathogenesis of a wide variety of major adverse pregnancy outcomes, including preeclampsia [6], ectopic pregnancy [7], intrauterine growth restriction [8], and recurrent miscarriage [9]. Moreover, sphingolipids, especially ceramides, have been identified in a number of cross-gestational studies to be associated with early-onset preeclampsia in pregnancy women during the first trimester, which make them candidate markers for early prediction of preeclampsia and other pregnancy-related disorders [10-11]. Therefore, in order to provide further insights into various conditions of pregnancy, we undertook a population-based study to establish the physiological baseline levels of ceramides and their biosynthetic precursors, dihydroceramides, during normal pregnancies.

To date, various analytical methodologies have been established for the measurement of ceramides in clinical specimens, including thin layer chromatography [12], normal phase liquid chromatography [13], immunochemistry [14], and gas chromatography [15]. However, these methods, in general, are limited by their narrow dynamic range, poor detection sensitivity, low analytic throughput, and inability to resolve individual ceramides at the molecular level, which preclude expansion of their application to routine clinical analysis. Recently, the advent of high-performance liquid chromatography interfaced to tandem mass spectrometer (MS/MS) has given rise to the development of ceramide assays with greater sensitivity, specificity, and throughput from complex biological matrices. Previously, Jiang et al. established a liquid chromatography-tandem mass spectrometric (LC/MS/MS) method for simultaneous quantification of ceramides (d18:1/22:0) and (d18:1/24:0) in human plasma [16], and Kauhanen et al. developed a high-throughput LC/MS/MS approach for routine clinical measurement of ceramides (d18:1/16:0), (d18:1/18:0), (d18:1/24:1), and (d18:1/24:0) in human plasma [17]. Both methods demonstrated good accuracy, precision, and throughput over clinically relevant ranges, but lack sufficient molecular coverage for the assay. Therefore, the expansion of the analytical capacity to include more ceramides and dihydroceramides in the assay would be useful.

In this study, we developed and validated a high-throughput LC/MS/MS method for parallel quantification of 16 ceramides and 10 dihydroceramides in human serum within 5 mins by using stable isotope-labeled ceramides as internal standards. This method employs a protein precipitation method for high-throughput sample preparation, reverse-phase isocratic elusion for chromatographic separation, and Multiple Reaction Monitoring (MRM) for mass spectrometric detection. The assay was validated against US FDA guidelines for LLOQ, linearity, precision, accuracy, extraction recovery, stability, and carryover. The validated assay was then applied to determine serological baselines of ceramides and dihydroceramides in cross-gestational normal pregnancies.

## 2. Materials and Methods

### 2.1 **Materials**

All calibration standards including ceramides (d18:1/14:0), (d18:1/16:0), (d18:1/17:0), (d18:1/18:1), (d18:1/18:0), (d18:1/20:0), (d18:1/22:0), (d18:1/24:1), (d18:1/24:0), and dihydroceramides (d18:0/16:0), (d18:0/18:1), (d18:0/18:0), (d18:0/24:1) were purchased from Avanti Lipids (Alabaster, AL). Stable isotope-labeled internal standards including d_7_-ceramide (d18:1/16:0), (d18:1/18:0), (d18:1/24:1), and (d18:1/24:0) were also purchased from Avanti Lipids. HPLC grade water, methanol, 2-propanol, and chloroform were obtained from Fisher Scientific (Pittsburgh, PA). Analytical grade ammonium bicarbonate was purchased from Sigma Aldrich (St. Louis, MO). The de-lipidized serum VD-DDC Mass Spec Gold was obtained from Golden West Biological (Temecula, CA). All materials were directly used without further purification.

### 2.2 Human Serum Sample

Serum samples from 10 healthy human donors were purchased from the Stanford Blood Center (Palo Alto, CA). These sample were combined to generate a pooled serum sample for quality control purposes.

Clinical samples containing 29 maternal serum samples from full-term pregnancies without complication were purchased from ProMedDX Inc. (Norton, MA, USA, http://www.promeddx.com) and included detailed case report forms. ProMedDX Inc. confirmed that all of these specimens were collected under Institutional Review Board-approved protocols by qualified Investigator sites. These sites conducted ProMedDX studies according to 21 CFR, ICH/GCP guidelines and HIPAA Privacy Regulations. Informed consent was obtained from every subject unless this requirement had been waived by the respective IRB.

All serum samples were aliquoted and stored at -80 °C prior to analysis.

### 2.3 Stock and Working Solutions

Stock solutions of ceramides (d18:1/14:0), (d18:1/16:0), (d18:1/17:0), (d18:1/18:1), (d18:1/18:0), (d18:1/20:0), (d18:1/22:0), (d18:1/24:1), (d18:1/24:0) and dihydroceramides (d18:0/16:0), (d18:0/18:1), (d18:0/18:0), (d18:0/24:1) were prepared by dissolving the lyophilized powders in MeOH:CHCl_3_ (1:1) to obtain a concentration of 5.00 mM.

A set of six-level calibrator working solutions were prepared by mixing and serially diluting stock solutions with 2-propanol to obtain concentrations at 1.00, 2.00, 10.0, 50.0, 2.00×10^2^, 1.00×10^3^ nM for ceramides (d18:1/14:0), (d18:1/17:0), (d18:1/18:1), (d18:1/18:0), (d18:1/20:0) and dihydroceramides (d18:0/16:0), (d18:0/18:1), (d18:0/18:0), (d18:0/24:1), and at 5.00, 10.0, 50.0, 2.50×10^2^, 1.00×10^3^, 5.00×10^3^ nM for ceramides (d18:1/16:0), (d18:1/22:0), (d18:1/24:1), (d18:1/24:0).

A set of four-level quality control (QC) working solutions were prepared by mixing and serially diluting stock solutions with 2-propanol to obtain concentrations at 1.00, 2.50, 1.00×10^2^, 7.50×10^2^ nM for ceramides (d18:1/14:0), (d18:1/17:0), (d18:1/18:1), (d18:1/18:0), (d18:1/20:0) and dihydroceramides (d18:0/16:0), (d18:0/18:1), (d18:0/18:0), (d18:0/24:1), and at 5.00, 12.5, 1.00×10^2^, 3.75×10^3^ for ceramides (d18:1/16:0), (d18:1/22:0), (d18:1/24:1), (d18:1/24:0).

The stock solutions of d_7_-ceramide (d18:1/16:0), (d18:1/18:0), (d18:1/24:1), and (d18:1/24:0) were prepared by dissolving the lyophilized powders in MeOH:CHCl_3_ (1:1) to obtain a universal concentration at 1.00 mM. The internal standard (IS) working solution was prepared by mixing and serially diluting stock solutions with methanol to obtain concentrations at 5.00 nM for all stable isotope-labeled ceramides.

The QC serum sample was prepared as described in Section 2.2 “Human Serum Sample” All prepared solutions and samples were stored at -20 °C prior to use.

### 2.4 Sample Preparation

For the preparation of blank samples, a 10-μL aliquot of de-lipidized serum was spiked with 10 μL of 2-propanol. The blank samples were extracted with 200 μL of methanol and the IS working solution to obtain double and single blanks, respectively.

For the preparation of calibrators, a 10-μL aliquot of de-lipidized serum was spiked with 10 μL of calibrator working solution at the corresponding level. The spiked calibrators were individually extracted with 200 μL of the IS working solution to obtain a set of calibrators based on 6 concentration levels.

For the preparation of QC samples, a 10-μL aliquot of de-lipidized serum was spiked with 10 μL of the QC working solution at the corresponding level. The spiked QC samples were individually extracted with 200 μL of the IS working solution to obtain a set of 4 concentration levels.

For the preparation of serum samples, a 10-μL aliquot of the unknown sample was spiked with 10 μL of 2-propanol and extracted with 200 μL of the IS working solution.

Following the extraction, all extracted samples were subjected to vigorous vortex for 30 secs and high-speed centrifuge at 12,000×g under 4 °C for 5 mins. Thereafter, 180 μL of supernatant was removed from each sample and transferred into an auto-sampler vial with micro-insert for LC/MS analysis.

### 2.5. LC/MS Instrumentation

The Dionex Ultimate 3000 UHPLC system consisted of a degasser, a RS binary pump, a RS auto-sampler, and a RS column compartment from Thermo Fisher (San Jose, CA). The UHPLC system was interfaced with a TSQ Quantiva mass spectrometer equipped with electrospray ionization source and a built-in Rheodyne switch valve from Thermo Fisher. Data acquisition and chromatographic peak integration were implemented using the XCalibur 4.0 software package from Thermo Fisher.

### 2.6. LC-MS/MS Procedure

Following sample preparation, 10 μL of the sample was injected onto an ACE Excel SuperC18 column (1.7 μm, 100 mm×2.1 mm; MAC-MOD Analytical, Chadds, PA). The mobile was composed of a mixture of methanol and 2-propanol at 1:1 buffered by 10 mM of ammonium bicarbonate. Chromatographic separation was carried out using a 5-min isocratic elusion program. Briefly, the LC eluent was directed to the waste for the first 1.0 min and then switched back to the electrospray interface from 1.1 to 5 min, allowing the targeted ceramides and dihydroceramides to be sequentially eluted, ionized, and detected by the system. The flow rate was set constantly at 0.3 mL/min, and temperatures of the auto-sampler and column oven were maintained at 4 and 30 °C, respectively, throughout the analysis.

The mass spectrometer was operated in a scheduled multiple reaction monitoring (MRM) mode to continuously acquire data from the LC eluent. The retention time-dependent data acquisition was employed using pre-defined retention time windows with variable widths (1.2 mins for medium chain and 1.5 mins for long chain ceramides and dihydroceramides) to record the extracted ion chromatograms (EIC) of targeted analytes. The MRM transitions for targeted ceramides and dihydroceramides were individually optimized by direct syringe pump infusion of 0.5 μM of the corresponding standards at 10 μL/min into the mass spectrometer in the presence of 10 mM of ammonium bicarbonate. The optimized MRM transitions with scheduled retention times are given in Table 1 and Table 2 for absolutely quantitated analytes, approximately quantitated analytes, and stable isotope-labeled internal standards. The Q1 and Q3 resolutions were both set at 0.7 Da, and the cycle time was set at 1 sec.

**Table 1.** The optimized MRM transitions for absolutely quantitated analytes

| **Compound** | **Product Ion** | **RT**<br>**(min)** | **RT**<br>**Window**<br>**(min)** | **Polarity** | **Precursor**<br>**(m/z)** | **Product**<br>**(m/z)** | **Collision**<br>**Energy**<br>**(V)** | **RF**<br>**Lens**<br>**(V)** |
| --- | --- | --- | --- | --- | --- | --- | --- | --- |
| Ceramide<br>(d18:1/14:0) | Quantitative | 1.58 | 1.20 | Negative | 508.5 | 252.3 | 27.19 | 121.0 |
|  | Qualitative | 1.58 | 1.20 | Negative | 508.5 | 226.2 | 25.78 | 121.0 |
| Ceramide<br>(d18:1/16:0) | Quantitative | 1.79 | 1.20 | Negative | 536.6 | 280.4 | 30.23 | 121.0 |
|  | Qualitative | 1.79 | 1.20 | Negative | 536.6 | 254.3 | 26.94 | 121.0 |
| Dihydroceramide<br>(d18:0/16:0) | Quantitative | 1.89 | 1.20 | Negative | 538.6 | 280.4 | 30.38 | 136.0 |
|  | Qualitative | 1.89 | 1.20 | Negative | 538.6 | 237.3 | 35.08 | 136.0 |
| Ceramide<br>(d18:1/17:0) | Qualitative | 1.91 | 1.20 | Negative | 550.6 | 294.4 | 28.96 | 101.0 |
|  | Quantitative | 1.91 | 1.20 | Negative | 550.6 | 268.3 | 27.60 | 101.0 |
| Ceramide<br>(d18:1/18:1) | Qualitative | 1.82 | 1.20 | Negative | 562.6 | 306.4 | 29.62 | 115.0 |
|  | Quantitative | 1.82 | 1.20 | Negative | 562.6 | 280.3 | 26.53 | 115.0 |
| Dihydroceramide<br>(d18:0/18:1) | Qualitative | 2.04 | 1.20 | Negative | 564.6 | 306.4 | 30.88 | 123.0 |
|  | Quantitative | 2.04 | 1.20 | Negative | 564.6 | 263.3 | 30.58 | 123.0 |
| Ceramide<br>(d18:1/18:0) | Qualitative | 2.04 | 1.20 | Negative | 564.6 | 308.4 | 31.29 | 129.0 |
|  | Quantitative | 2.04 | 1.20 | Negative | 564.6 | 282.3 | 28.56 | 129.0 |
| Dihydroceramide<br>(d18:0/18:0) | Quantitative | 2.18 | 1.20 | Negative | 566.6 | 308.4 | 31.44 | 140.0 |
|  | Qualitative | 2.18 | 1.20 | Negative | 566.6 | 265.3 | 35.94 | 140.0 |
| Ceramide<br>(d18:1/20:0) | Quantitative | 2.36 | 1.20 | Negative | 592.6 | 336.4 | 32.40 | 136.0 |
|  | Qualitative | 2.36 | 1.20 | Negative | 592.6 | 310.3 | 29.97 | 136.0 |
| Ceramide<br>(d18:1/22:0) | Quantitative | 2.74 | 1.20 | Negative | 620.7 | 364.5 | 34.52 | 132.0 |
|  | Qualitative | 2.74 | 1.20 | Negative | 620.7 | 338.4 | 32.05 | 132.0 |
| Ceramide<br>(d18:1/24:1) | Quantitative | 2.75 | 1.50 | Negative | 646.7 | 390.5 | 36.19 | 142.0 |
|  | Qualitative | 2.75 | 1.50 | Negative | 646.7 | 364.4 | 30.83 | 142.0 |
| Dihydroceramide<br>(d18:0/24:1) | Quantitative | 3.25 | 1.50 | Negative | 648.7 | 390.5 | 35.33 | 138.0 |
|  | Qualitative | 3.25 | 1.50 | Negative | 648.7 | 347.4 | 35.33 | 138.0 |
| Ceramide<br>(d18:1/24:0) | Quantitative | 3.25 | 1.50 | Negative | 648.7 | 392.5 | 34.62 | 145.0 |
|  | Qualitative | 3.25 | 1.50 | Negative | 648.7 | 366.4 | 31.99 | 145.0 |

**Table 2.** The optimized MRM transitions for approximately quantitated analytes and stable isotope-labeled internal standards

| **Compound** | **Product Ion** | **RT**<br>**(min)** | **RT**<br>**Window**<br>**(min)** | **Polarity** | **Precursor**<br>**(m/z)** | **Product**<br>**(m/z)** | **Collision**<br>**Energy**<br>**(V)** | **RF**<br>**Lens**<br>**(V)** |
| --- | --- | --- | --- | --- | --- | --- | --- | --- |
| Ceramide<br>(d18:1/20:1) | Quantitative | 2.06 | 1.20 | Negative | 590.6 | 334.4 | 32.40 | 136.0 |
|  | Qualitative | 2.06 | 1.20 | Negative | 590.6 | 308.3 | 29.97 | 136.0 |
| Dihydroceramide<br>(d18:0/20:0) | Quantitative | 2.52 | 1.20 | Negative | 594.6 | 336.4 | 31.44 | 140.0 |
|  | Qualitative | 2.52 | 1.20 | Negative | 594.6 | 293.3 | 35.94 | 140.0 |
| Ceramide<br>(d18:1/21:0) | Quantitative | 2.54 | 1.20 | Negative | 606.6 | 350.4 | 32.40 | 136.0 |
|  | Qualitative | 2.54 | 1.20 | Negative | 606.6 | 324.3 | 29.97 | 136.0 |
| Dihydroceramide<br>(d18:0/21:0) | Quantitative | 2.72 | 1.20 | Negative | 608.6 | 350.4 | 31.44 | 140.0 |
|  | Qualitative | 2.72 | 1.20 | Negative | 608.6 | 307.3 | 35.94 | 140.0 |
| Ceramide<br>(d18:1/22:1) | Quantitative | 2.38 | 1.20 | Negative | 618.7 | 362.5 | 34.52 | 132.0 |
|  | Qualitative | 2.38 | 1.20 | Negative | 618.7 | 336.4 | 32.05 | 132.0 |
| Dihydroceramide<br>(d18:0/22:0) | Quantitative | 2.94 | 1.20 | Negative | 622.7 | 364.5 | 35.38 | 133.0 |
|  | Qualitative | 2.94 | 1.20 | Negative | 622.7 | 321.4 | 35.33 | 133.0 |
| Ceramide<br>(d18:1/23:0) | Quantitative | 2.99 | 1.50 | Negative | 634.7 | 378.5 | 34.52 | 132.0 |
|  | Qualitative | 2.99 | 1.50 | Negative | 634.7 | 352.4 | 32.05 | 132.0 |
| Dihydroceramide<br>(d18:0/23:0) | Quantitative | 3.22 | 1.50 | Negative | 636.7 | 378.5 | 35.38 | 133.0 |
|  | Qualitative | 3.22 | 1.50 | Negative | 636.7 | 335.4 | 35.33 | 133.0 |
| Ceramide<br>(d18:1/24:2) | Quantitative | 2.43 | 1.50 | Negative | 644.7 | 388.5 | 36.19 | 142.0 |
|  | Qualitative | 2.43 | 1.50 | Negative | 644.7 | 362.4 | 30.83 | 142.0 |
| Ceramide<br>(d18:1/25:0) | Quantitative | 3.50 | 1.50 | Negative | 662.7 | 406.5 | 34.62 | 145.0 |
|  | Qualitative | 3.50 | 1.50 | Negative | 662.7 | 380.4 | 31.99 | 145.0 |
| Dihydroceramide<br>(d18:0/25:0) | Quantitative | 3.80 | 1.50 | Negative | 664.7 | 406.5 | 35.33 | 138.0 |
|  | Qualitative | 3.80 | 1.50 | Negative | 664.7 | 363.4 | 35.33 | 138.0 |
| Ceramide<br>(d18:1/26:0) | Quantitative | 3.85 | 1.50 | Negative | 676.7 | 420.5 | 34.62 | 145.0 |
|  | Qualitative | 3.85 | 1.50 | Negative | 676.7 | 394.4 | 31.99 | 145.0 |
| Dihydroceramide<br>(d18:0/26:0) | Quantitative | 4.15 | 1.50 | Negative | 678.7 | 420.5 | 35.33 | 138.0 |
|  | Qualitative | 4.15 | 1.50 | Negative | 678.7 | 377.4 | 35.33 | 138.0 |
| d<sub>7</sub>-Ceramide<br>(d18:1/16:0) | Quantitative | 1.79 | 1.20 | Negative | 543.6 | 280.4 | 30.23 | 121.0 |
| d<sub>7</sub>-Ceramide<br>(d18:1/18:0) | Quantitative | 2.04 | 1.20 | Negative | 571.6 | 308.4 | 31.29 | 129.0 |
| d<sub>7</sub>-Ceramide<br>(d18:1/24:1) | Quantitative | 2.72 | 1.50 | Negative | 653.7 | 390.5 | 36.19 | 142.0 |
| d<sub>7</sub>-Ceramide<br>(d18:1/24:0) | Quantitative | 3.22 | 1.50 | Negative | 655.7 | 392.5 | 34.62 | 145.0 |

In addition, the source parameters were also optimized by in-source mixing of the mobile phase flow at 0.3 mL/min with continuous infusion of 0.5 μM of standard cocktail at 10 μL/min via a tee connector. The spray voltage was optimized at 3500 V, and the optimal gas flows were determined to be 30, 10, and 0 Arb for Sheath Gas, Aux Gas, and Sweep Gas, respectively. The temperatures for ion transfer tube and vaporizer were also optimized to be both at 300 °C.

### 2.7 Qualification

The qualification of targeted ceramides and dihydroceramides was implemented via a two-step process. In the first step, the chromatographic peak areas of both quantitative and qualitative ions were integrated for absolutely quantitated analytes from Table 1 based on their retention times in corresponding EICs across all calibrators, QCs, and serum samples. The integrated peak areas were first manually inspected, normalized by their labeled counterparts/analogs, and then calculated for product ion ratios by dividing the area under curve (AUC) of quantitative ions by the AUC of qualitative ions. In order for the absolutely quantitated analytes to be accepted for the subsequent quantitation process, they must be qualified to show the mean product ion ratio deviation within the 10% tolerance cutoff in the serum samples relative to the calibrator and QC samples.

In the second step, the chromatographic peak areas of both quantitative and qualitative ions were integrated for approximately quantitated analytes from Table 2 based on their observed retention times in corresponding EICs across all serum samples. The integrated peak areas were first manually inspected, normalized by their labeled counterparts/analogs, and then calculated for product ion ratios by dividing the AUC of quantitative ions by the AUC of qualitative ions. In consideration of the unavailability of commercial standards for the approximately quantitated analytes, a 20% acceptance cutoff was set on the mean product ion ratio deviations in serum samples relative to their absolutely quantitated analogs in calibrator and QC samples for qualification based on a pre-defined relationship described in Table 3.

**Table 3.**
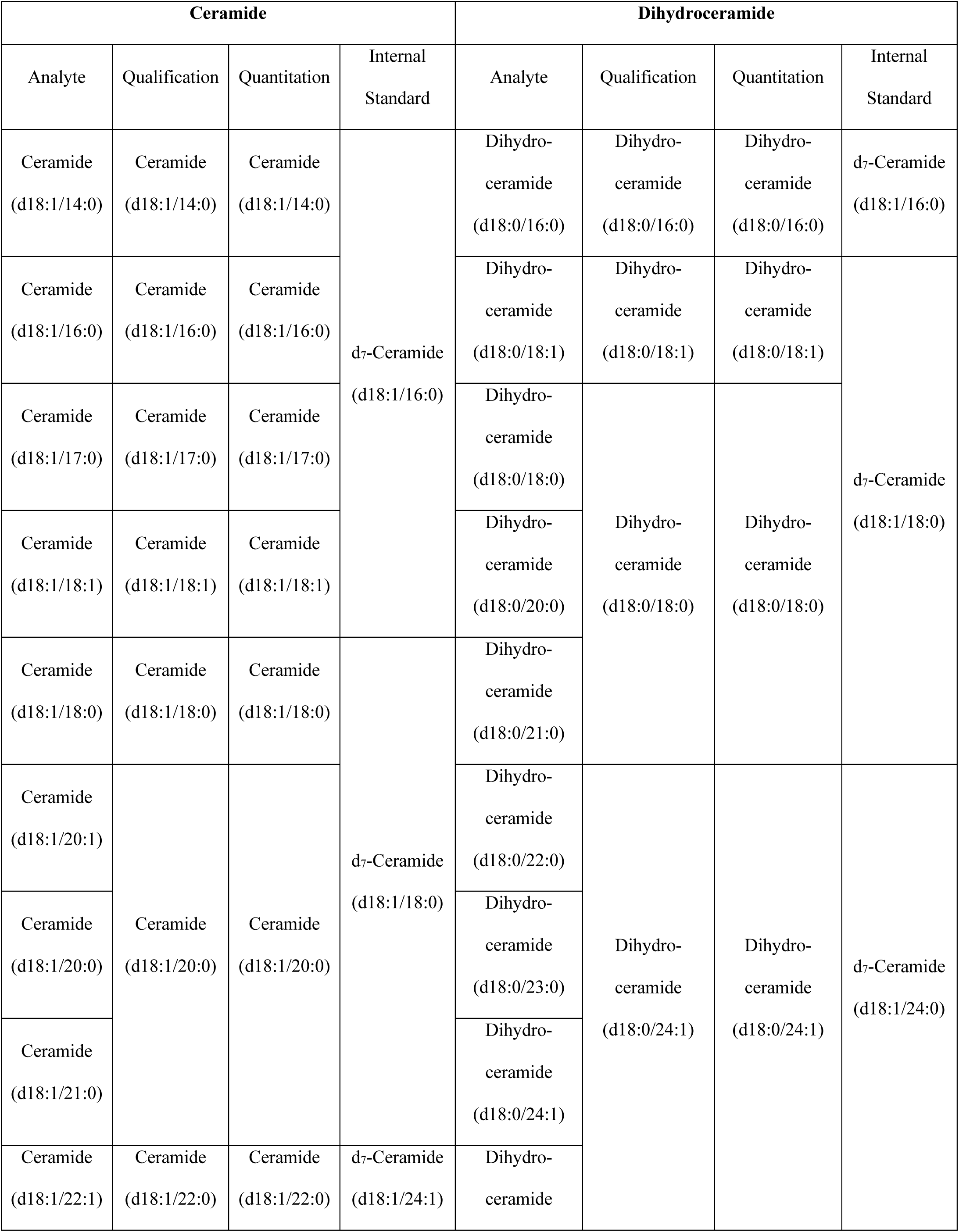
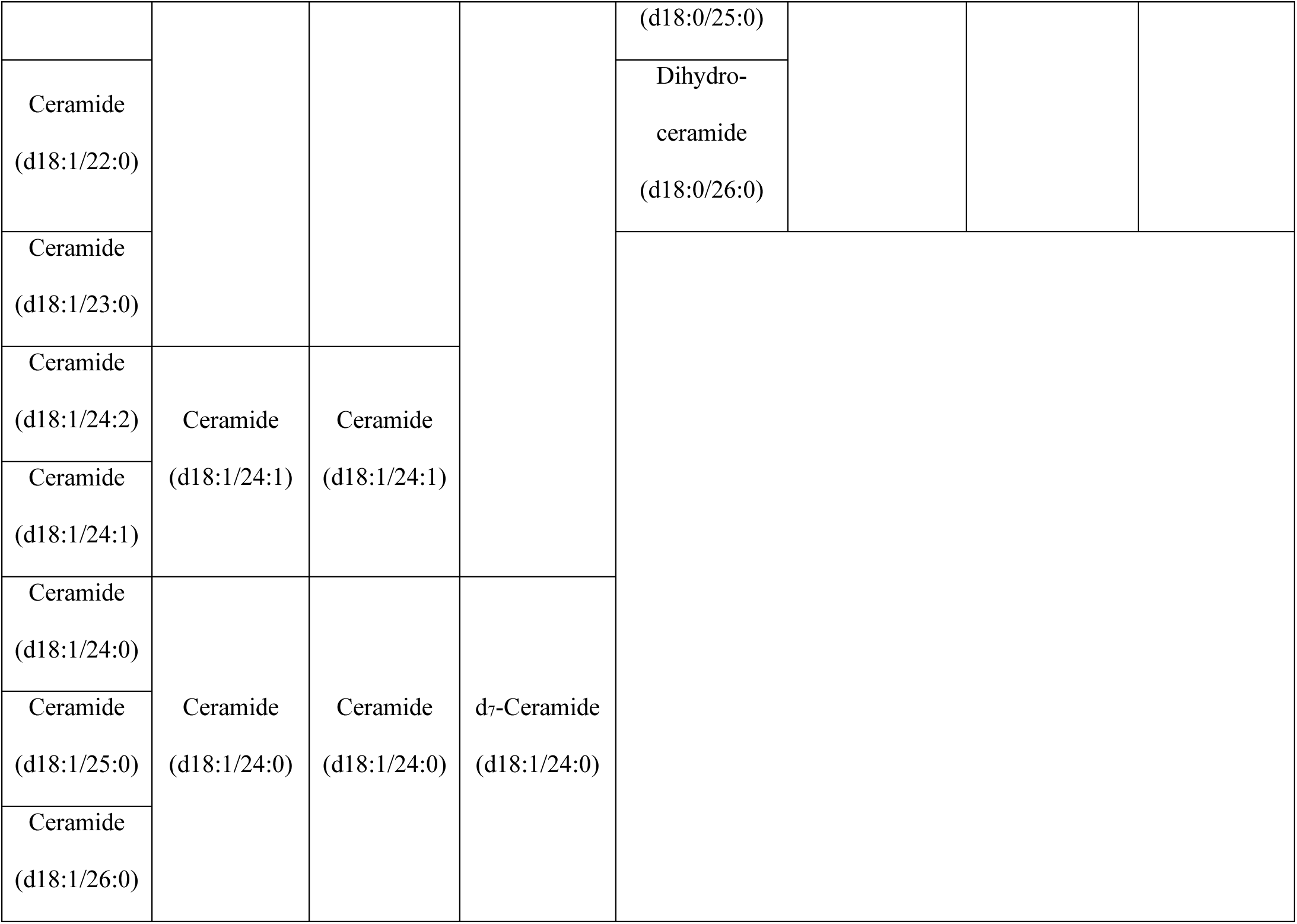
The relationships of normalization, qualification, and calibration for quantitation

The normalization assignment as well as the structural homology relationship are detailed in Table 3.

### 2.8 Quantitation

The quantitation of targeted ceramides and dihydroceramides was implemented via two separate approaches. In the first approach, known as absolute quantitation, the IS-normalized peak area ratios for absolutely quantitated analytes from Table 1 were plotted against the spiked concentrations in calibrators to establish calibration curves based on 6 levels. Linear regression fitting with a weighting factor of 1/x^2^ was employed for the calibration. Thereafter, a >0.99 cutoff was set on the square of the correlation coefficient to ensure that individual calibration curves were qualified for quantitation. After that, the IS-normalized peak area ratios of targeted analytes were plugged into the corresponding calibration curves to obtain absolutely quantitated concentrations in blank, QC, and serum samples.

In the second approach, known as approximate quantitation, the IS-normalized peak area ratios for approximately quantitated analytes from Table 2 were plugged into the assigned calibration curves based on the extent of structural homology to obtain approximately quantitated concentrations in the serum samples.

The analyte-specific assignment for calibration and quantitation is detailed in Table 2.

### 2.9 Statistic Analysis

For the clinical portion of the study, quadratic polynomial fitting was used to generate gestation-dependent serological baselines for targeted ceramides and dihydroceramides in normal pregnancies. Student’s t-test was applied to determine the significance of differences between different pregnancy groups.

## Result and Discussion

### 3.1 **MS Condition**

Electrospray ionization interfaced with tandem mass spectrometer (ESI-MS/MS) appears to be a more advantageous platform than the immuno-based approach for analysis of ceramides, considering its superior selectivity to unambiguously differentiate molecular ceramides with major structural resemblance, which would be technically challenging for the immune-based method due to the significant inter-ceramide cross-reactivity observed with the antibodies [18]. The ionization mechanisms and in-depth structural characterization of collision-induced dissociation (CID) fragments have been well documented for ceramides in ESI-MS/MS [19-23]. In ESI+ mode, the ionization of ceramides generally takes place at the carbonyl oxygen from the C2 amide linkage via the addition of a single proton, which, upon dissociation, generates a characteristic fragment at 264.3 m/z, corresponding to the protonated d18:1 sphingosine ion after the loss of two water molecules. This signature fragment represents the essential d18:1 sphingosine building block, which forms the principal infrastructure for both qualitative and quantitative analysis of ceramides by ESI+. However, due to the fragile nature of hydroxy groups, significant in-source dehydration from protonated molecular ions has been observed on the ceramides during the ionization in ESI+, leading to reduced abundance of molecular ions and thus diminished detection sensitivity [19-21]. In ESI-mode, the ceramide undergoes ionization by removing a proton from the C2 amide nitrogen and, upon dissociation, produces a wide spectrum of structurally informative fragments, featured by the neutral loss (NL) of 256.2 m/z, corresponding to the loss of a hexadecenal and water molecules. The fragment generated by NL of 256.2 m/z in ESI-has been demonstrated to be a superior fragment over the 264.3 m/z fragment in ESI+ for both qualitative and quantitative analysis, as it is not limited by the detrimental in-source dehydration phenomenon observed during the protonation. However, the deprotonation of ceramides in ESI-could be severely affected by the presence of chloride ion, a direct competitor for the uncharged ceramide molecule during ionization, significantly dropping the abundance of the deprotonated molecular ion by forming the chloride adduct [22-23]. In this study, ESI-was selected as the method for ionization over ESI+ based on the following rationales: 1) once dissociated, the molecular ion of ceramide in ESI-produces more structurally informative fragments than ESI+, allowing the unambiguous identification and quantification of ceramides in extremely complex biological matrices; 2) the signal-diluting effect imposed by the chloride ion on ionization in ESI-could be practically overcome by separating the sample across a hydrophobic stationary phase to deplete the chloride ion and enrich the molecular ceramides prior to ionization; 3) elimination of the in-source dehydration phenomenon in the negative ionization efficiently preserves the deprotonated molecular ion of ceramide to the maximal extent, therefore boosting detection sensitivity.

The representative product ion spectra are shown in Fig. 2 for ceramide (d18:1/18:0) and dihydroceramide (d18:0/18:0). As illustrated by their MS/MS fingerprints, ceramide (d18:1/18:0) could be readily identified by the presence of N-vinyloctadecanamide and octadecanamide anions generated by the NL of 256.2 and 282.3 m/z from the molecular ion, whereas dihydroceramide (d18:0/18:0), which differs by one double bond, was characterized by the presence of N-vinyloctadecenamide and hexadecylketene anions produced by the NL of 258.2 and 301.3 m/z from the molecular ion. In addition, the fragments created by the NL of 256.2 and 258.2 m/z were found to be the most abundant product ions for ceramides and dihydroceramides, respectively. These observations were consistent with the previous finding [23].

**Fig.2.**
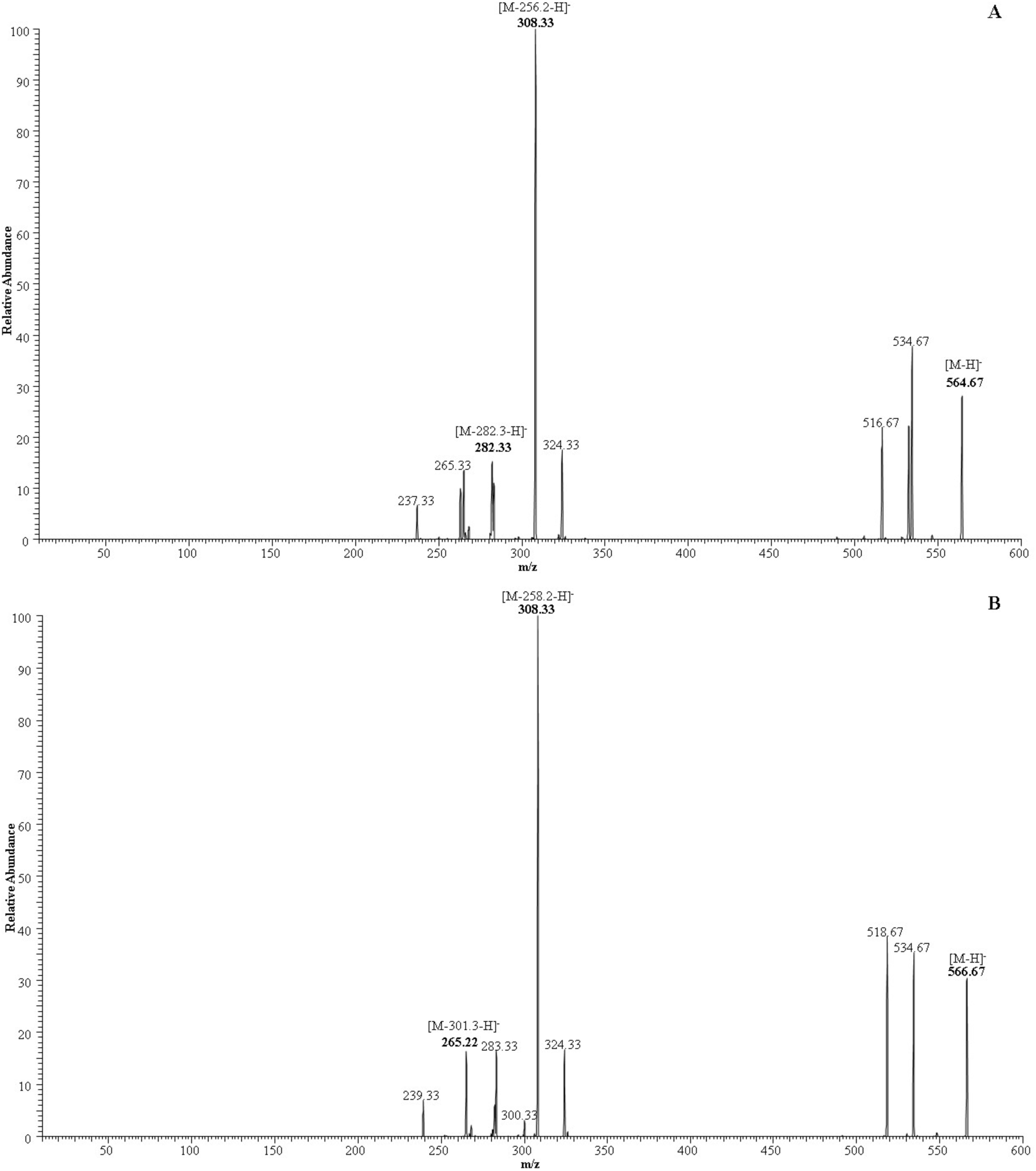
Fragmentation spectra of ceramide (d18:1/18:0) (A) and dihydroceramide (d18:0/18:0) (B).

To assure the sensitivity and specificity of the MS detection, multiple fragments were selected as product ion candidates and paired with [M-H]^-^ as the precursor ion to be evaluated in terms of absolute abundance, fragmentation reproducibility, and robustness to interference for the targeted analysis by MRM. As a result, the product ions [M-H-256.2]^-^ and [M-H-282.3]^-^ from ceramides were observed to outweigh others for quantitative and qualitative analysis, respectively. Likewise, the product ions [M-H-258.2]^-^ and [M-H-301.3]^-^ from dihydroceramides were identified to be superior over others for quantitative and qualitative analysis, respectively. The optimized SRM transitions are listed in Table 1 and 2. The fragmentation mechanisms of 256.2 and 282.3 m/z NL for ceramides and 258.2 and 301.3 m/z NL for dihydroceramides are shown in Fig. 3.

**Fig.3.**
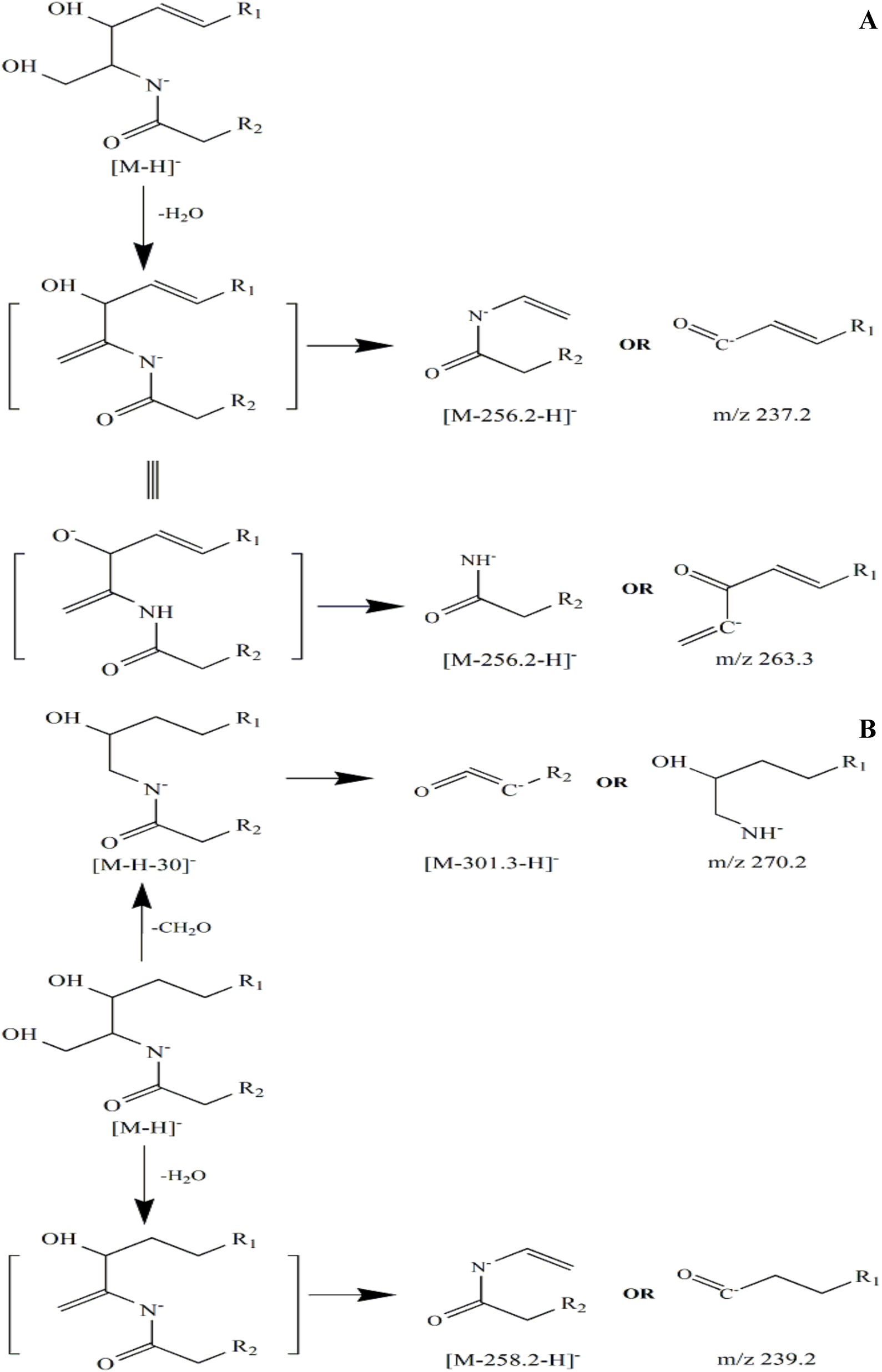
Fragmentation mechanism of NL 256.2 and 282.3 m/z for ceramide (A) and NL 258.2 and 301.3 m/z for dihydroceramide (B).

**Fig. 4.**
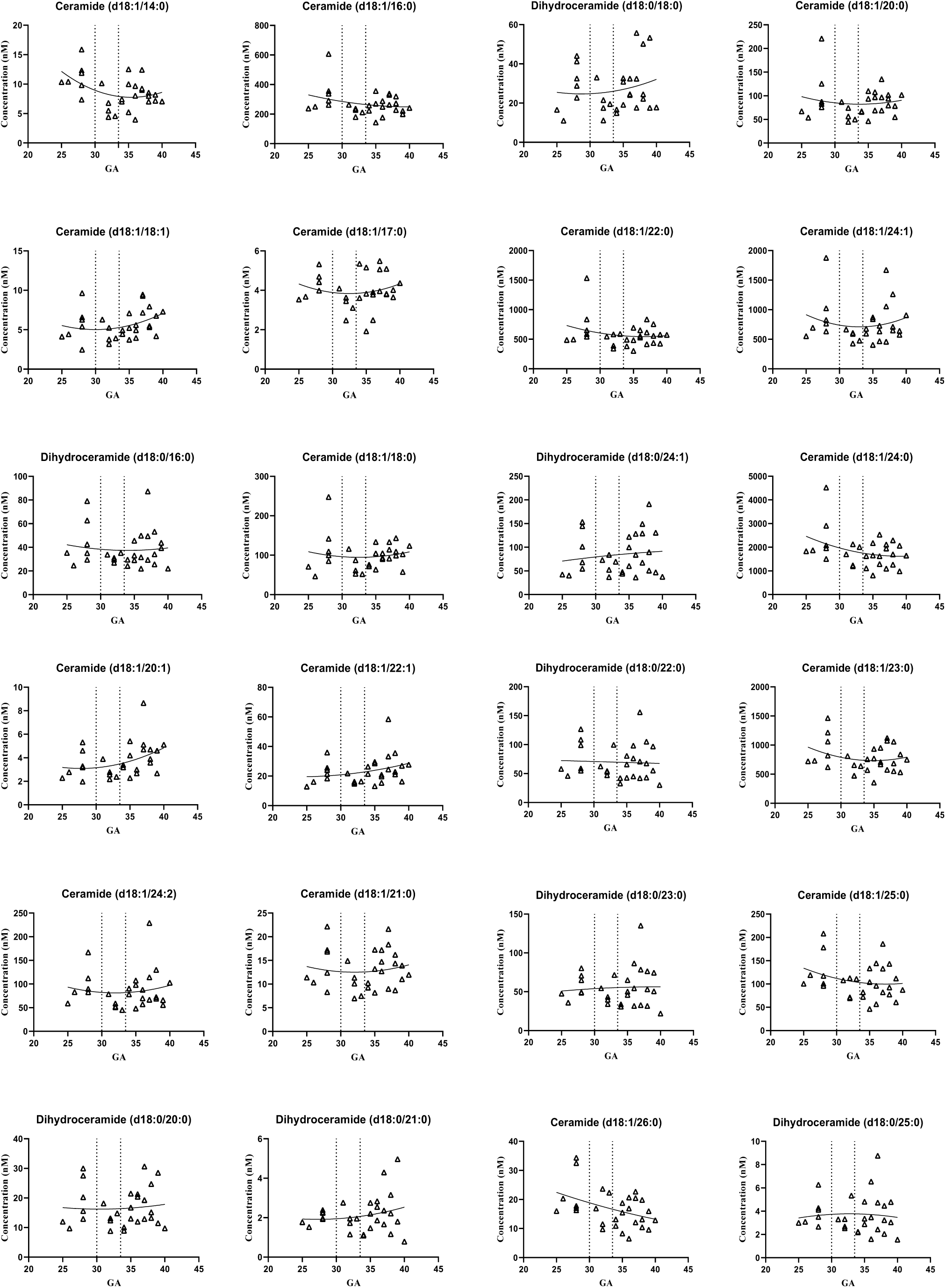
Gestational age-dependent serological baseline plots of individual ceramides and dihydroceramides from normal pregnancy samples. The X-axis represents the gestational age in weeks, and the Y-axis represents the concentrations of the analyte in nM. The two dotted lines represent the 30 weeks and 34 weeks of gestational ages.

**Fig.5.**
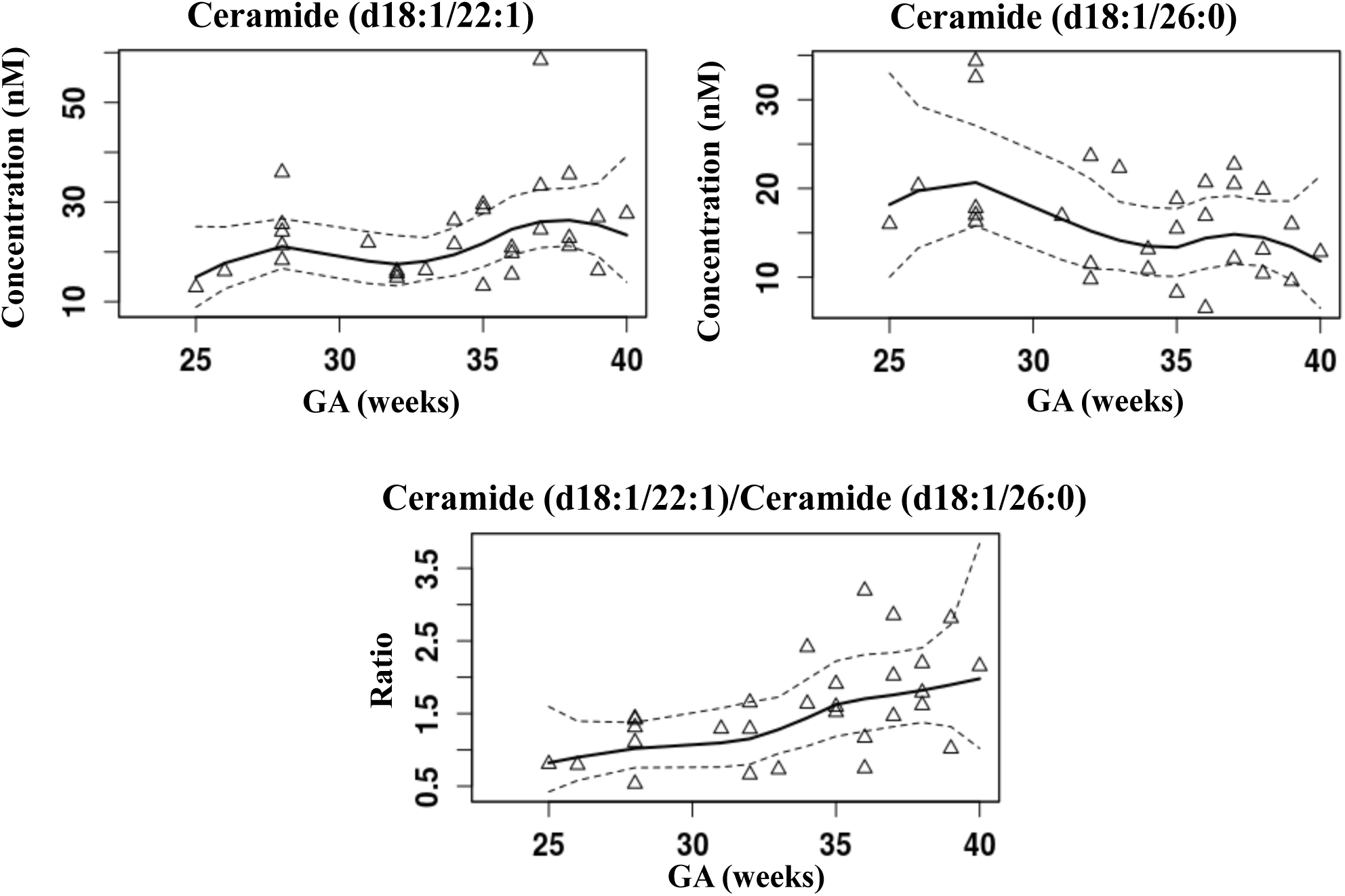
Gestational age-dependent serological baselines of ceramide (d18:1/22:1), (d18:1/26:0), and their ratio from normal pregnancy samples with the 95% confidence interval defined by the dotted lines.

### 3.2 LC Condition

According to previous studies [24-27], a prior chromatographic separation step is essential to alleviate the charging competition that occurs between co-eluting analytes and matrix components and minimize the extent of ionization suppression introduced onto the targeted analytes, thus improving the sensitivity, specificity, and robustness of the assay. Among multiple chromatographic platforms, reverse phase chromatography is known for its capability to retain and separate lipophilic molecules in order of increasing hydrophobicity, and its application has been documented in multiple studies to be fundamental for the chromatographic separation of glycosphingolipids [24-25], lipophilic vitamins [26], and steroids [27] to take place in a highly reproducible manner. As a result, in view of the hydrophobic nature of ceramides and dihydroceramides as complex sphingolipids, an ACE Excel SuperC18 UPLC column (1.7 μm, 100 mm×2.1 mm) was selected for chromatographic separation to provide reliable matrix cleanup and analyte enrichment prior to the ionization.

Moreover, in the existing methodologies, gradient elusion programs were preferentially employed for the analysis of ceramides to improve chromatographic resolution and detection sensitivity. Based on published data, utilization of gradient elusion programs allows the targeted ceramides to be retained, separated, and eluted reproducibly with excellent run-to-run consistency [24-27]. However, as the tradeoff of improved chromatographic resolution and detection sensitivity, gradient elusion is also technically subjected to a number of limitations. On one hand, the continuously variable mobile phase percentage program used by gradient elusion unfavorably influences the stability of the chromatographic baseline, causing the baseline to fluctuate constantly throughout the analysis. On the other hand, gradient elusion inevitably requires additional chromatographic periods for column cleanup and re-equilibration, leading to a shortened analyte elusion window, prolonged running time, and extended instrument idle period. As an alternative, isocratic elusion seems to be a potential solution to circumvent these shortcomings. In practice, isocratic elusion renders a more stabilized chromatographic baseline with minimal column cleanup and re-equilibration steps required. Unfortunately, isocratic elusion also suffers from pitfalls such as a broadened chromatographic peak, a tendency to column overloading, and enhanced sample carryover. Therefore, in the course of method development, a number of vital chromatographic parameters including mobile phase composition, sample loading solvent, column oven temperature, flow rate, sample injection volume, and running time were individually and collectively optimized to obtain improved chromatographic resolution with sharpened peaks in the absence of column overloading. The sample carryover was also evaluated under optimized conditions to be negligible from sample to sample. It is worth noting that with the use of methanol, both as extraction solvent and sample loading solvent, the width of the chromatographic peak was significantly narrowed to allow for increased signal intensity and a lowered detection limit. In addition, the inclusion of 2-propanol in the mobile phase minimizes the buildup of hydrophobic matrix components onto the column over time and improves the elution of targeted analytes with sharpened chromatographic peaks and a shortened analytical run. Under our optimized conditions, the targeted ceramides and dihydroceramides were consistently eluted off the column from 1.5 to 4.5 min in an order of increased hydrophobicity across a 3-min retention time window.

Chemically, the pKa value for the amide N-H bond on ceramide should range from 25 to 26, suggesting that the amide nitrogen is essentially a weak acid and very unlikely to donate its proton under acidic or neutral pH. However, in the presence of a base in large excess, the acid-base interaction would be expected to facilitate the proton-donating process on the amide N-H bond, stabilize the amide nitrogen anion that is generated, and thus increase ionization efficiency. Furthermore, in consideration of the subsequent detection phase, the base being used must be a volatile chemical that is compatible with the ESI interface and MS analysis, ranking the ammonium bicarbonate buffer the top candidate on the list. In addition, as the majority of C18 reverse phase columns are unable to be operated under basic pH owing to the chemical hydrolysis that might occur between the alkyl stationary phase and the silica bed, the ACE Excel SuperC18 UPLC column we selected for the analysis was further confirmed by the manufacturer to possess a stable alkyl stationary phase under extended pH range from 2 to 11, allowing the high pH chromatographic separation to be carried out with promising robustness.

### 3.3 Approximate Quantitation

Approximate quantitation, known as the approximate determination of certain molecules by referencing their signal responses to the calibration of resembled structural analogs established in the same matrix, has been used as an alternative method to circumvent the obstacles pertaining to conventional calibration using a one-to-one matched standard, and to provide approximately quantitative insights into molecules with unusual structures in the absence of commercial standards. The application of this approximate quantitation approach has been exemplified by multiple studies for the analysis of phosphatidylcholines [28], hexosylceramides [29], phosphatidylethanolamines [30], sulfatides [31], triglycerides [32], and cholesteryl esters [33]. In those studies, lipid standards with unnatural fatty acid moieties but identical lipid head groups to the analytes were selected as class-specific representatives to build the concentration-dependent calibration in the studied samples, which is then used as the known references to obtain quantitative information from the targeted analytes within the same lipid class by approximate quantitation via mathematic approach. In view of the distinctive structural diversity of lipids, the utilization of exogenous analogs as class-specific representative calibrators has been demonstrated to analytically bypass the interfering signals originating from the endogenous lipids, simplifying the overall calibration process, and thus allowing the simultaneous quantitation of hundreds of lipid species from the sample in a rapid and reproducible fashion, paving the way for the shotgun lipidomic analysis. A number of prerequisites were proposed to be essential for the qualification of the approximate quantitation approach: 1) the molecular moiety that is subjected to structural variation should have minimal influence on the ionization efficiency, 2) the representative analogs selected for calibration should be structurally homologous to the targeted analytes, and 3) the linearity of representative calibrators should be demonstrated a priori in the matched matrix across the relevant concentration ranges. As mentioned above, for ceramides and dihydroceramides, the deprotonation process for negative ionization is predominantly undertaken at the nitrogen proton from the amide-linkage, and the variation on aliphatic chain length and saturation degree supposedly accounts for minimal impact on ionization efficiency. In addition, the linearities of representative ceramides and dihydroceramides with greatest structural similarities to the approximately quantitated analytes were established in the matched matrix over the meaningful concentration ranges, making the approximate quantitation approach as the most favorable strategy for measurement of those unusual ceramides and dihydroceramides in the clinical samples. However, as the complexity of the matrix components from human serum samples raises concerns about the validity of the detected signal, especially on the approximately quantitated analytes, approximately quantitative results obtained from clinical samples were required to undergo a series of qualification procedures against the pre-defined acceptance criteria to ensure the selectivity of the measurement.

### 3.4 Method Validation

The developed method was validated against the bioanalytical method validation guideline published by the US FDA in 2018 to evaluate for LLOQ, linearity, precision, accuracy, extraction recovery, stability, and carryover [34].

### 3.4.1 LLOQ

The LLOQ, defined as the concentration level with S/N ≥10, for each absolutely quantitated analyte was determined by spiking the corresponding unlabeled standard at known concentration into the de-lipidized serum and then serially diluting the spiked serum sample with de-lipidized serum until S/N=10 was reached in six replicates. In addition, the double blank and single blank samples were also analyzed in parallel to ensure that the baseline signal intensity from presented interference consistently accounted for <10% of the signal intensity at the LLOQ for each absolutely quantitated analyte. The determined LLOQ levels are given in Table 4.

**Table 4.** The LLOQ, linear range, and linearity of individual calibration curves

| **Analyte** | **LLOQ**<br>**(nM)** | **Linear Range**<br>**(nM)** | **Weighting**<br>**Factor** | **R<sup>2</sup>** | **Calibration Curve Equation** |
| --- | --- | --- | --- | --- | --- |
| Ceramide<br>(d18:1/14:0) | 1.00 | 1.00-1.00×10<sup>3</sup> | 1/X<sup>2</sup> | 0.999 | Y = -0.00022648+0.0092549*X |
| Ceramide<br>(d18:1/16:0) | 5.00 | 5.00-5.00×10<sup>3</sup> | 1/X<sup>2</sup> | 0.999 | Y = 0.00904432+0.0116833*X |
| Ceramide<br>(d18:1/17:0) | 1.00 | 1.00-1.00×10<sup>3</sup> | 1/X<sup>2</sup> | 0.999 | Y = 0.00051912+0.0128024*X |
| Ceramide<br>(d18:1/18:1) | 1.00 | 1.00-1.00×10<sup>3</sup> | 1/X<sup>2</sup> | 0.999 | Y = 0.00003149+0.0148810*X |
| Ceramide<br>(d18:1/18:0) | 1.00 | 1.00-1.00×10<sup>3</sup> | 1/X<sup>2</sup> | 0.999 | Y = 0.00149927+0.0100025*X |
| Ceramide<br>(d18:1/20:0) | 1.00 | 1.00-1.00×10<sup>3</sup> | 1/X<sup>2</sup> | 1.000 | Y = 0.00057816+0.0107459*X |
| Ceramide<br>(d18:1/22:0) | 5.00 | 5.00-5.00×10<sup>3</sup> | 1/X<sup>2</sup> | 0.999 | Y = -0.00134115+0.0080520*X |
| Ceramide<br>(d18:1/24:1) | 5.00 | 5.00-5.00×10<sup>3</sup> | 1/X<sup>2</sup> | 0.999 | Y = 0.00489232+0.0087511*X |
| Ceramide<br>(d18:1/24:0) | 5.00 | 5.00-5.00×10<sup>3</sup> | 1/X<sup>2</sup> | 0.999 | Y = 0.01361550+0.0075987*X |
| Dihydroceramide<br>(d18:0/16:0) | 1.00 | 1.00-1.00×10<sup>3</sup> | 1/X<sup>2</sup> | 1.000 | Y = 0.00464943+0.0163816*X |
| Dihydroceramide<br>(d18:0/18:1) | 1.00 | 1.00-1.00×10<sup>3</sup> | 1/X<sup>2</sup> | 0.999 | Y = 0.00016294+0.0132201*X |
| Dihydroceramide<br>(d18:0/18:0) | 1.00 | 1.00-1.00×10<sup>3</sup> | 1/X<sup>2</sup> | 0.999 | Y = 0.00215833+0.0125412*X |
| Dihydroceramide<br>(d18:0/24:1) | 1.00 | 1.00-1.00×10<sup>3</sup> | 1/X<sup>2</sup> | 0.999 | Y = -0.00045146+0.0113406*X |

### 3.4.2 Linearity

The linearity of the calibration curve, defined as the square of correlation coefficient (r^2^) of the linear regression curve, was determined for each absolutely quantitated analyte based on 6 concentration levels with a weighting factor of 1/x^2^. The linear ranges were from 1.00 to 1.00×10^3^ nM for ceramides (d18:1/14:0), (d18:1/17:0), (d18:1/18:1), (d18:1/18:0), (d18:1/22:0) and dihydroceramides (d18:0/16:0), (d18:0/18:1), (d18:0/18:0), (d18:0/24:1), and from 5.00 to 5.00×10^3^ nM for ceramides (d18:1/16:0), (d18:1/20:0), (d18:1/24:1), (d18:1/24:0). The r^2^ values for all calibration curves, which are presented in Table 4, were >0.99.

### 3.4.3 Precision and Accuracy

The precision, defined as the coefficient of variation (CV%) of multiple measurements based on multiple sampling of the same homogenous sample, and accuracy, defined as the percent error (PE%) of the determined value relative to the nominal value, were evaluated for intra- and inter-assay measurements by analyzing QC samples prepared at four different concentrations, namely LLOQ, Low, Medium, and High, in six replicates for four independent runs. As shown in Table 5, the CV% and PE% were <11.9 and 10.6% for the intra-assay measurement and <6.57 and 9.53% for inter-assay measurement. In addition, the QC pooled serum sample was prepared and analyzed in a similar setting to assess the precision of the approximately quantitated analytes for intra- and inter-assay measurements, since the pure unlabeled standards were not commercially available for those analytes. As shown in Table 6, the CV% values were <8.85 and 13.3% for intra- and inter-assay measurements, respectively, demonstrating the reproducibility of the assay.

**Table 5.** Intra- & inter-assay precision and accuracy for internal QC samples

| Analyte | QC Level | Nominal Concentration (nM) | Intra-Assay |  | Inter-Assay |  |
| --- | --- | --- | --- | --- | --- | --- |
|  |  |  | PE (%) | CV (%) | PE (%) | CV (%) |
| Ceramide (d18:1/14:0) | LLOQ | 1.00 | 10.6 | 2.96 | 3.78 | 4.43 |
|  | Low | 2.50 | 2.16 | 4.17 | 1.55 | 2.86 |
|  | Medium | 1.00×10<sup>2</sup> | 0.50 | 2.41 | 1.10 | 0.76 |
|  | High | 7.50×10<sup>2</sup> | 5.57 | 3.06 | 9.53 | 2.53 |
| Ceramide (d18:1/16:0) | LLOQ | 5.00 | 5.34 | 6.06 | 1.20 | 6.18 |
|  | Low | 12.5 | 1.51 | 4.08 | 0.27 | 1.86 |
|  | Medium | 1.00×10<sup>2</sup> | 1.62 | 2.35 | 3.55 | 1.48 |
|  | High | 3.75×10<sup>3</sup> | 0.87 | 2.99 | 3.11 | 2.69 |
| Ceramide (d18:1/17:0) | LLOQ | 1.00 | 1.85 | 6.22 | 4.55 | 2.81 |
|  | Low | 2.50 | 0.09 | 5.82 | 2.18 | 3.21 |
|  | Medium | 1.00×10<sup>2</sup> | 5.04 | 2.06 | 3.60 | 1.84 |
|  | High | 7.50×10<sup>2</sup> | 2.77 | 3.32 | 4.63 | 1.26 |
| Ceramide (d18:1/18:1) | LLOQ | 1.00 | 5.54 | 5.48 | 0.13 | 5.27 |
|  | Low | 2.50 | 0.57 | 3.91 | 0.43 | 1.86 |
|  | Medium | 1.00×10<sup>2</sup> | 3.33 | 2.38 | 3.71 | 2.58 |
|  | High | 7.50×10<sup>2</sup> | 2.17 | 2.31 | 5.30 | 2.54 |
| Ceramide (d18:1/18:0) | LLOQ | 1.00 | 0.22 | 3.79 | 2.52 | 6.88 |
|  | Low | 2.50 | 5.55 | 5.38 | 2.37 | 2.85 |
|  | Medium | 1.00×10<sup>2</sup> | 1.57 | 5.15 | 3.65 | 3.69 |
|  | High | 7.50×10<sup>2</sup> | 3.28 | 4.50 | 6.49 | 3.44 |
| Ceramide (d18:1/20:0) | LLOQ | 1.00 | 1.73 | 11.9 | 0.84 | 1.81 |
|  | Low | 2.50 | 0.43 | 1.88 | 0.43 | 6.97 |
| | Medium | $1.00 \times 10^2$ | 0.95 | 2.82 | 2.95 | 3.09 |
| | High | $7.50 \times 10^2$ | 4.41 | 2.71 | 6.44 | 1.78 |
| Ceramide<br>(d18:1/22:0) | LLOQ | 5.00 | 8.76 | 3.92 | 3.78 | 3.31 |
|  | Low | 12.5 | 1.28 | 5.64 | 1.67 | 3.19 |
| | Medium | $1.00 \times 10^2$ | 2.80 | 4.28 | 4.08 | 1.92 |
| | High | $3.75 \times 10^3$ | 0.40 | 4.03 | 1.51 | 2.36 |
| Ceramide<br>(d18:1/24:1) | LLOQ | 5.00 | 3.38 | 2.61 | 0.89 | 4.85 |
|  | Low | 12.5 | 0.93 | 4.68 | 1.62 | 2.20 |
| | Medium | $1.00 \times 10^2$ | 3.97 | 3.86 | 4.85 | 1.36 |
| | High | $3.75 \times 10^3$ | 0.33 | 4.72 | 2.40 | 2.84 |
| Ceramide<br>(d18:1/24:0) | LLOQ | 5.00 | 3.01 | 2.86 | 1.62 | 6.15 |
|  | Low | 12.5 | 2.35 | 5.45 | 2.36 | 3.19 |
| | Medium | $1.00 \times 10^2$ | 4.94 | 2.97 | 3.47 | 1.73 |
| | High | $3.75 \times 10^3$ | 0.06 | 4.54 | 1.39 | 2.77 |
| Dihydroceramide<br>(d18:0/16:0) | LLOQ | 1.00 | 5.52 | 14.6 | 2.65 | 3.19 |
|  | Low | 2.50 | 4.50 | 2.55 | 1.50 | 2.87 |
| | Medium | $1.00 \times 10^2$ | 1.05 | 3.79 | 2.13 | 1.79 |
| | High | $7.50 \times 10^2$ | 1.11 | 3.80 | 3.74 | 1.93 |
| Dihydroceramide<br>(d18:0/18:1) | LLOQ | 1.00 | 2.12 | 4.57 | 2.20 | 2.77 |
|  | Low | 2.50 | 0.45 | 6.56 | 0.50 | 1.13 |
| | Medium | $1.00 \times 10^2$ | 2.57 | 3.24 | 4.68 | 1.88 |
| | High | $7.50 \times 10^2$ | 0.17 | 3.81 | 1.96 | 1.48 |
| Dihydroceramide<br>(d18:0/18:0) | LLOQ | 1.00 | 4.33 | 3.13 | 2.99 | 6.88 |
|  | Low | 2.50 | 5.93 | 7.23 | 0.32 | 5.11 |
| | Medium | $1.00 \times 10^2$ | 3.68 | 3.55 | 3.18 | 0.61 |
| | High | $7.50 \times 10^2$ | 4.79 | 4.36 | 5.14 | 1.83 |
| Dihydroceramide<br>(d18:0/24:1) | LLOQ | 1.00 | 3.31 | 4.67 | 1.51 | 5.67 |
|  | Low | 2.50 | 1.70 | 6.08 | 1.28 | 0.94 |
| | Medium | $1.00\times10^2$ | 4.43 | 4.11 | 1.22 | 3.81 |
| | High | $7.50\times10^2$ | 4.54 | 3.14 | 3.05 | 5.25 |

**Table 6.**
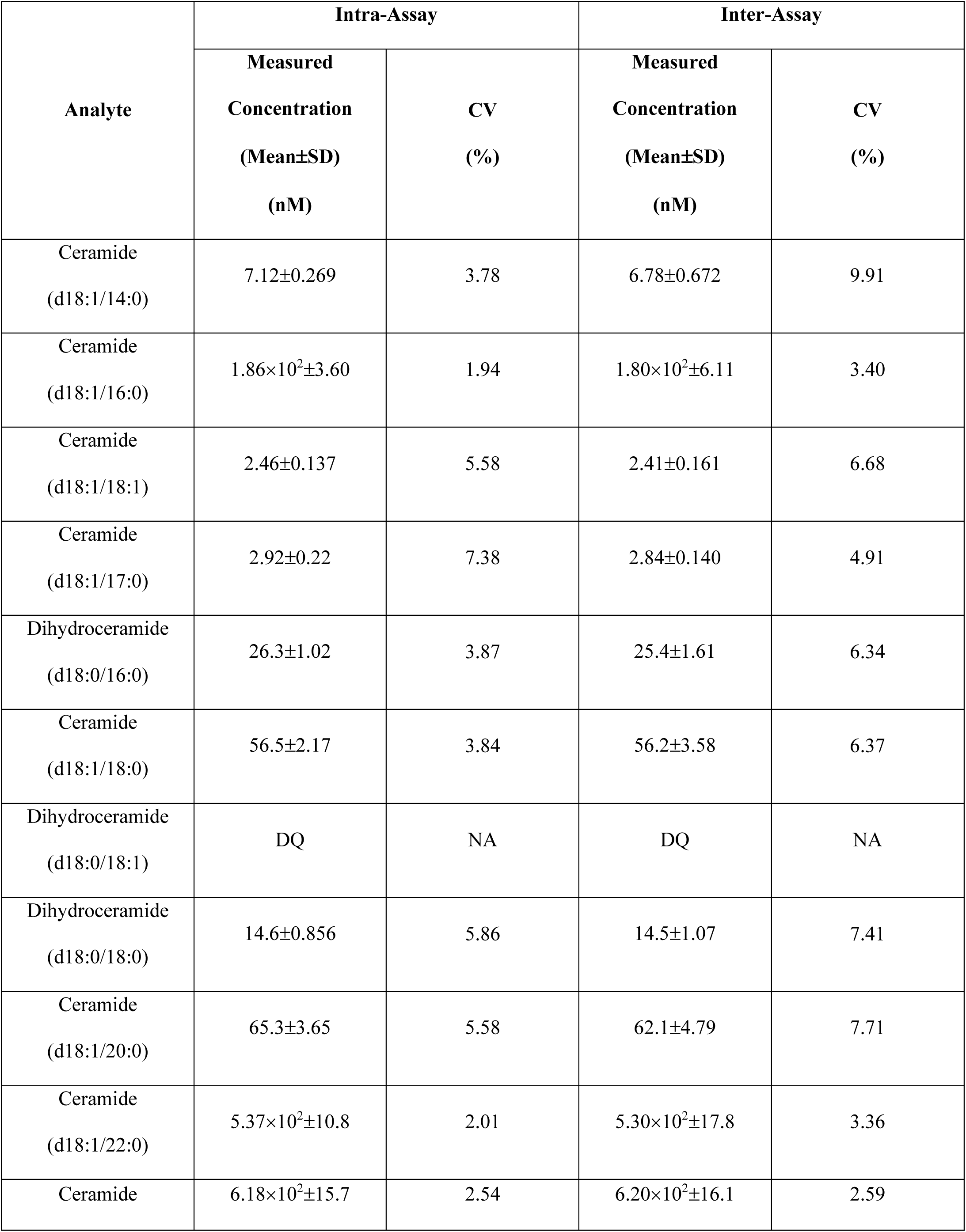
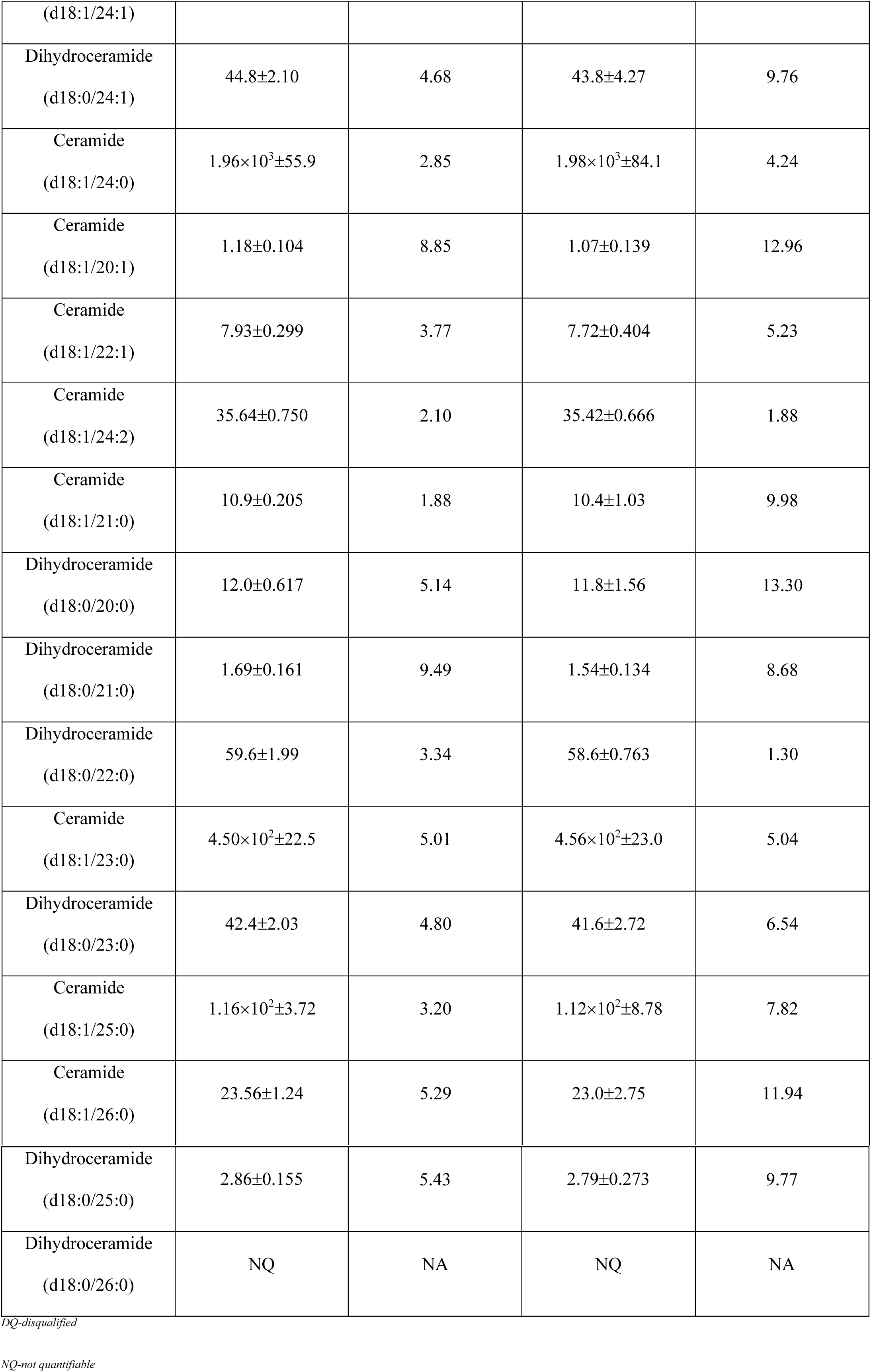
Intra- & Inter-assay precision for pooled serum sample

### 3.5.4 Extraction Recovery

Extraction recovery, defined as a percentage of the known amount of an analyte carried through the sample extraction and processing steps of the method, was evaluated for the extraction protocol by comparing two sets of QC samples prepared at three different concentrations, namely Low, Medium, and High, that undergo analyte spiking before and after the extraction in four replicates. As shown in Table 7, the percentage recovery values were >89.4% for all analytes at studied concentrations, indicating insignificant loss of the analytes during sample extraction and processing steps.

**Table 7.** Extraction recovery and stability for internal QC samples

| **Analyte** | **QC Level** | **Nominal Concentration (nM)** | **Extraction Recovery (Mean±SD) (%)** | **Autosampler Stability (Mean±SD) (%)** | **Benchtop Stability (Mean±SD) (%)** | **Long-term Stability (Mean±SD) (%)** |
| --- | --- | --- | --- | --- | --- | --- |
| Ceramide (d18:1/14:0) | Low | 2.50 | 91.1±3.3 | 98.3±2.7 | 96.6±0.5 | 83.2±4.9 |
|  | Medium | 1.00×10<sup>2</sup> | 93.3±3.9 | 104.3±2.6 | 99.4±2.0 | 89.8±1.3 |
|  | High | 7.50×10<sup>2</sup> | 89.4±5.5 | 99.3±1.9 | 96.5±3.0 | 87.2±1.9 |
| Ceramide (d18:1/16:0) | Low | 12.5 | 102.3±3.2 | 92.8±3.8 | 98.2±3.0 | 83.3±1.6 |
|  | Medium | 1.00×10<sup>2</sup> | 104.5±1.7 | 97.1±1.1 | 99.1±2.0 | 88.0±1.1 |
|  | High | 3.75×10<sup>3</sup> | 102.4±1.0 | 98.2±0.5 | 95.5±1.6 | 88.4±3.0 |
| Ceramide (d18:1/17:0) | Low | 2.50 | 92.7±0.6 | 100.3±3.4 | 96.6±1.6 | 82.0±1.3 |
|  | Medium | 1.00×10<sup>2</sup> | 95.1±2.0 | 99.2±5.1 | 96.0±4.4 | 84.3±0.7 |
|  | High | 7.50×10<sup>2</sup> | 105.6±0.8 | 99.2±2.2 | 97.0±2.0 | 90.1±1.6 |
| Ceramide (d18:1/18:1) | Low | 2.50 | 98.2±6.2 | 93.8±3.9 | 95.4±2.6 | 85.1±2.6 |
|  | Medium | 1.00×10<sup>2</sup> | 101.8±5.6 | 95.8±4.2 | 96.9±3.8 | 88.8±2.4 |
|  | High | 7.50×10<sup>2</sup> | 106.3±8.0 | 98.5±0.3 | 95.2±1.4 | 88.7±1.0 |
| Ceramide (d18:1/18:0) | Low | 2.50 | 97.3±6.2 | 94.8±4.6 | 99.3±5.9 | 86.7±2.0 |
|  | Medium | 1.00×10<sup>2</sup> | 97.2±4.4 | 94.7±4.3 | 96.8±0.9 | 86.4±1.7 |
|  | High | 7.50×10<sup>2</sup> | 99.2±2.2 | 99.9±3.8 | 97.1±0.9 | 89.7±2.1 |
| Ceramide (d18:1/20:0) | Low | 2.50 | 100.9±5.0 | 93.1±3.2 | 95.9±5.6 | 84.7±4.8 |
|  | Medium | 1.00×10<sup>2</sup> | 99.4±0.4 | 99.4±3.7 | 96.5±3.6 | 87.1±1.3 |
|  | High | 7.50×10<sup>2</sup> | 95.8±3.2 | 99.8±2.6 | 96.2±6.7 | 88.8±1.8 |
| Ceramide (d18:1/22:0) | Low | 12.5 | 100.2±3.7 | 95.7±1.5 | 96.5±3.3 | 90.2±4.3 |
|  | Medium | 1.00×10<sup>2</sup> | 98.9±0.5 | 93.8±1.9 | 95.7±1.8 | 89.1±1.4 |
| | High | $3.75 \times 10^3$ | $101.7 \pm 1.6$ | $96.2 \pm 0.8$ | $98.8 \pm 4.9$ | $88.3 \pm 0.9$ |
| Ceramide<br>(d18:1/24:1) | Low | 12.5 | $100.5 \pm 3.8$ | $92.5 \pm 4.0$ | $94.8 \pm 3.5$ | $87.7 \pm 1.9$ |
| | Medium | $1.00 \times 10^2$ | $98.5 \pm 2.1$ | $92.3 \pm 0.9$ | $96.8 \pm 0.6$ | $87.9 \pm 1.7$ |
| | High | $3.75 \times 10^3$ | $103.2 \pm 0.3$ | $98.1 \pm 3.1$ | $96.6 \pm 3.3$ | $88.4 \pm 2.8$ |
| Ceramide<br>(d18:1/24:0) | Low | 12.5 | $100.2 \pm 2.4$ | $97.9 \pm 2.0$ | $101.5 \pm 3.8$ | $90.4 \pm 8.8$ |
| | Medium | $1.00 \times 10^2$ | $99.6 \pm 1.0$ | $94.9 \pm 1.3$ | $97.7 \pm 1.2$ | $86.9 \pm 1.0$ |
| | High | $3.75 \times 10^3$ | $100.0 \pm 0.2$ | $94.6 \pm 2.4$ | $93.3 \pm 1.0$ | $87.5 \pm 3.5$ |
| Dihydroceramide<br>(d18:0/16:0) | Low | 2.50 | $90.1 \pm 4.2$ | $104.4 \pm 3.5$ | $98.9 \pm 4.5$ | $90.7 \pm 2.1$ |
| | Medium | $1.00 \times 10^2$ | $103.7 \pm 7.8$ | $101.3 \pm 3.7$ | $101.2 \pm 3.0$ | $90.4 \pm 1.6$ |
| | High | $7.50 \times 10^2$ | $107.2 \pm 0.6$ | $100.5 \pm 2.3$ | $94.8 \pm 3.4$ | $90.3 \pm 3.0$ |
| Dihydroceramide<br>(d18:0/18:1) | Low | 2.50 | $101.8 \pm 1.0$ | $103.5 \pm 1.6$ | $100.6 \pm 3.2$ | $85.4 \pm 5.3$ |
| | Medium | $1.00 \times 10^2$ | $97.2 \pm 5.2$ | $101.2 \pm 5.6$ | $97.4 \pm 3.7$ | $89.6 \pm 1.4$ |
| | High | $7.50 \times 10^2$ | $97.8 \pm 5.9$ | $102.8 \pm 1.4$ | $96.3 \pm 4.6$ | $90.1 \pm 0.6$ |
| Dihydroceramide<br>(d18:0/18:0) | Low | 2.50 | $104.5 \pm 8.8$ | $100.0 \pm 6.2$ | $97.6 \pm 1.1$ | $85.3 \pm 3.0$ |
| | Medium | $1.00 \times 10^2$ | $101.2 \pm 2.9$ | $99.2 \pm 5.4$ | $96.0 \pm 2.4$ | $85.5 \pm 4.0$ |
| | High | $7.50 \times 10^2$ | $99.2 \pm 6.8$ | $102.0 \pm 1.0$ | $96.5 \pm 6.0$ | $88.4 \pm 2.7$ |
| Dihydroceramide<br>(d18:0/24:1) | Low | 2.50 | $100.5 \pm 3.8$ | $96.2 \pm 1.0$ | $97.9 \pm 5.2$ | $92.2 \pm 8.0$ |
| | Medium | $1.00 \times 10^2$ | $98.5 \pm 2.1$ | $94.8 \pm 1.6$ | $97.4 \pm 0.9$ | $89.6 \pm 2.6$ |
| | High | $7.50 \times 10^2$ | $103.2 \pm 0.3$ | $97.1 \pm 2.5$ | $99.8 \pm 4.2$ | $92.9 \pm 1.5$ |

### 3.4.5 Stability

Stability, defined as the percentage of intact analyte in a given matrix under specific storage and use conditions relative to the starting amount, for a given interval, was evaluated for auto-sampler, benchtop, and long-term storage scenarios by analyzing four sets of QC samples prepared at three different concentrations, namely Low, Medium, and High, that undergo storage at specific conditions. The storage conditions were 24 hours at 4°C, 4 hours at room temperature, and 21 days at -20°C for auto-sampler, benchtop, and long-term stabilities, respectively. As shown in Table 7, the stability percentages were >83.2% for all analytes under all conditions, suggesting minimal loss of analytes during storage.

### 3.4.6 Carryover

Carryover, defined as the appearance in a sample of an analyte from the proceeding sample, was evaluated by analyzing the calibrator sample with the highest analyte concentrations followed by the injection of a single blank sample with no analytes presented. The carryover effect was determined to be <0.01% for all analytes by including a needle washing step both before and after the injection of methanol.

### 3.5 Method Application

The ceramides and dihydroceramides in sera from 29 normal pregnancies were measured using the validated method. The determined concentrations are summarized in Table 8. The mean concentrations of ceramides (d18:1/16:0), (d18:1/18:0), and (d18:1/24:1) over all samples were 2.70×10^2^, 0.99×10^2^, and 7.69×10^2^ nM, respectively, which were in line with the clinical reference ranges reported by Mayo Clinic (1.90∼3.60×10^2^, 0.50∼1.40×10^2^, and 0.65∼1.65×10^3^ nM for ceramides (d18:1/16:0), (d18:1/18:0), and (d18:1/24:1), respectively). The concentrations of the ceramides and dihydroceramides as a function of GA are shown in Fig. 4. Monotonic changes along with the GA were observed in 13 analytes (7 increased and 6 decreased). Among the 13 analytes, ceramide (d18:1/22:1) increased most rapidly and ceramide (d18:1/26:0) decreased most rapidly. The ratio of two ceramides demonstrated a significant level change from 24-29 weeks to 34-40 weeks (p-value=7.43×10^-4^), indicating a possible physiological activity regulated by ceramides from mid- to late- gestation.

**Table 8.**
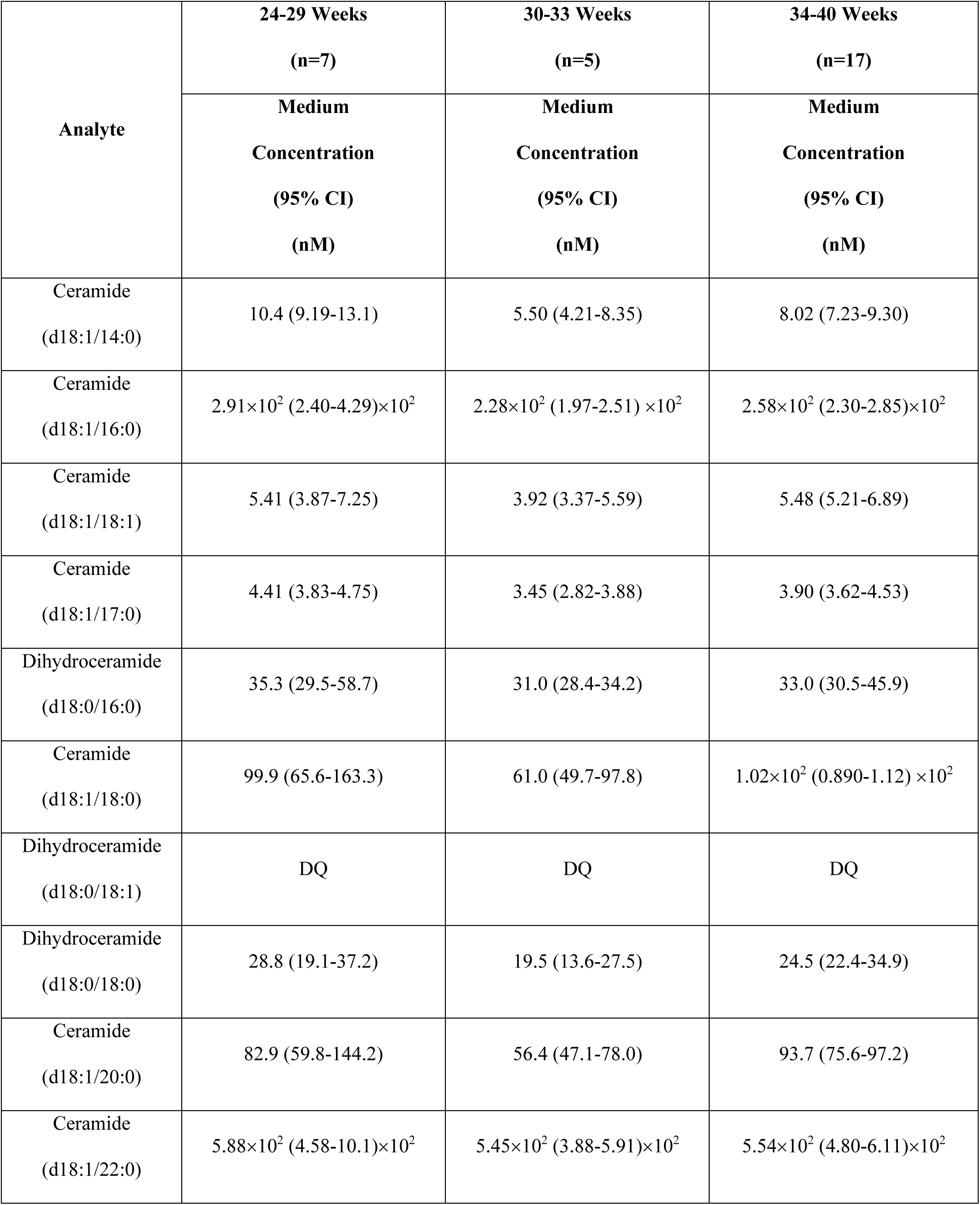
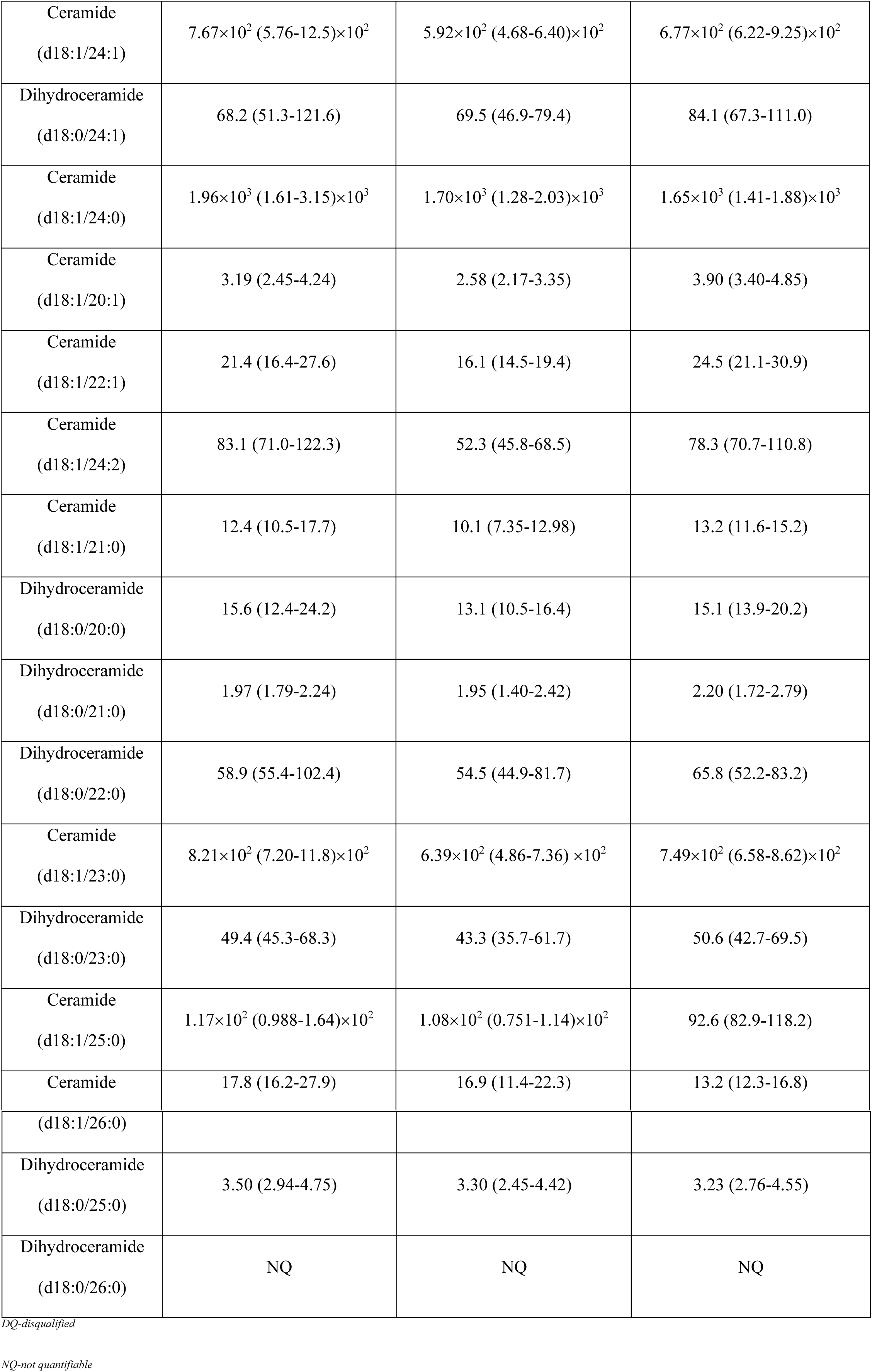
Levels of ceramides and dihydroceramides from measurement of application study

Our study, for the first time, characterized 8 endogenous ceramide and dihydroceramide species with novel chemical structures and quantified their levels in human blood, with high sensitivity and specificity accomplished by our new methodology. The new species were ceramides (d18:1/17:0), (d18:1/20:1), (d18:1/21:0), (d18:1/24:2) and dihydroceramides (d18:0/18:1), (d18:0/21:0), (d18:0/23:0), (d18:0/25:0). The majority of these unusual analytes were odd-chain ceramides and dihydroceramides derived from odd-chain fatty acids, which are not endogenously produced by the human body. Dairy products and meat from ruminant animals have been suggested to be important sources of odd-chain saturated fatty acids [35]. In addition, population-based studies have identified an inverse relationship between plasma levels of odd-chain saturated fatty acids and the risk of coronary heart disease [36] and type 2 diabetes [37], suggesting the potentially distinctive role of low abundant odd-chain ceramides in relation to the disease etiology and pathogenesis.

Ceramides, as metabolic messengers, are crucial to cell membrane stability and are able to act as bioactive lipids in cell signaling pathways, including immune responses, inflammation, cell cycle arrest, and apoptosis [38], and their modulation in pregnancy could serve as potential molecular targets to comprehensively understand the underlying pathophysiology of related disorders such as preeclampsia and preterm birth. Dihydroceramides, known as transiently produced intermediates during the de novo ceramide synthesis, have previously been deemed to be biologically inert [39], but recent evidence has suggested that they exert distinct biological functions that are complementary to the prevalent ceramides in a variety of disorders, including autophagy and hypoxia [39-41]. However, further research is required to better understand their pathophysiological association with the pregnancy disorder.

### 4. Conclusion

This study presents the development and validation of a high-throughput LC/MS/MS method for simultaneous quantification of 16 ceramides and 10 dihydroceramides in human serum within 5 mins by using stable isotope-labeled ceramides as internal standards. Our method employs a simple protein precipitation method for sample preparation, reverse phase isocratic elusion for chromatographic separation, and Multiple Reaction Monitoring (MRM) for mass spectrometric detection. The validated assay was utilized to determine serological baselines of ceramides and dihydroceramides in normal pregnancy across gestation. A ceramide ratio (d18:1/22:1 to d18:1/26:0) with a unique pattern associated with specific gestational ages was observed. In view of its high sensitivity, specificity, throughput, and low volume sample requirement, this method is expected to provide quantitative insights into the biology of sphingolipids during normal pregnancy, making it a potentially good practical approach to monitor the health status of pregnancy progression.

## Availability of data and materials

The datasets used and/or analyzed in this study are available upon request to the corresponding author.

## Competing interests

The authors declare that they have no competing interests.

## LIST OF ABBREVIATIONS

LC/MS/MS: Liquid chromatography-tandem mass spectrometric
IS: Internal standard
MRM: Multiple Reaction Monitoring
LLOQ: Lower Limit of Quantitation
ER: Endoplasmic reticulum
DEGS1: Dihydroceramide desaturase
MS/MS: Tandem mass spectrometer
QC: Quality control
EIC: Extracted ion chromatograms
AUC: Area under curve
ESI: Electrospray ionization
CID: Collision-induced dissociation
NL: Neutral loss
CV: Coefficient of variation
PE: Percent error
GA: Gestational age

## Acknowledgement

We thank colleagues at March of the Dimes Prematurity Research Center and Pediatrics Proteomics Group for critical discussions.

